# Temporal genomic imprinting regulates grain filling in barley

**DOI:** 10.64898/2026.11.04.764152

**Authors:** Martin Kovacik, Adam Lampar, Anna Nowicka, Marta Peirats-Llobet, Peter Šály, Kay Trafford, Zuzana Korchanová, Kateřina Kaduchová, Beáta Strejčková, Magnus Wohlfahrt Rasmussen, Søren Knudsen, Aparna Sreeprakash, Marie Červenková, Tereza Dobisová, Christoph Dockter, James Whelan, Clément Lafon Placette, Ales Pecinka

## Abstract

In cereals, the persistent endosperm accumulates starch and storage proteins, making their grains a major source of food and feed^1–3^. Early grain development is epigenetically controlled through the parent-of-origin-specific expression of imprinted genes in the endosperm that regulate endosperm proliferation, nutrient accumulation and embryo nourishment^4–6^. Imprinted genes have been identified in several major cereals, and a few have been functionally characterized^7–14^. However, it is unclear how imprinted genes regulate the accumulation of storage compounds in cereal grains. Through spatiotemporal transcriptome analysis of 255 imprinted genes in barley (*Hordeum vulgare*), we identify endosperm transfer cells in grains as a hotspot of expression of imprinted genes. Furthermore, we show that the transcription factor *LYS3* dynamically regulates grain filling in barley through early expression of the maternal allele, followed by partial activation of the paternal allele.

**One-sentence summary:** We identified imprinted genes in developing barley endosperm and characterized transcription factor *LYS3* as a temporally maternally expressed gene, uniparentally regulating barley grain filling.

## Results and Discussion

To identify imprinted genes in barley at the genome-wide level, we sequenced endosperm transcriptome at 8 and 12 days after pollination (DAP), corresponding to cellularizing and cellularized stages, respectively, from two different reciprocal F_1_ hybrid grains: (i) cultivar Morex (MOR) crossed with Ethiopian landrace (ETH), and (ii) MOR with wild barley (HS) (Fig. 1a). We identified 255 candidate imprinted genes, including 150 maternally expressed genes (MEGs) and 105 paternally expressed genes (PEGs) (Fig. 1b, c, Supplementary Fig. 1a, Supplementary Data Sets 1-5). The parent-of-origin specific expression was independently validated with highly consistent results (Supplementary Fig. 2). Substantial numbers of unique MEGs and PEGs in individual crosses were mostly due to hybrid-specific diagnostic SNPs (Supplementary Fig. 3). We observed similar numbers of imprinted genes between 8 and 12 DAP timepoints (104 versus 124 MEGs, 52.0% shared, and 87 versus 70 PEGs, 49.5% shared, respectively; Fig. 1c, Supplementary Data Set 5). Hence, we identified barley imprinted genes active predominantly during its endosperm cellularization and filling.

**Fig. 1.**
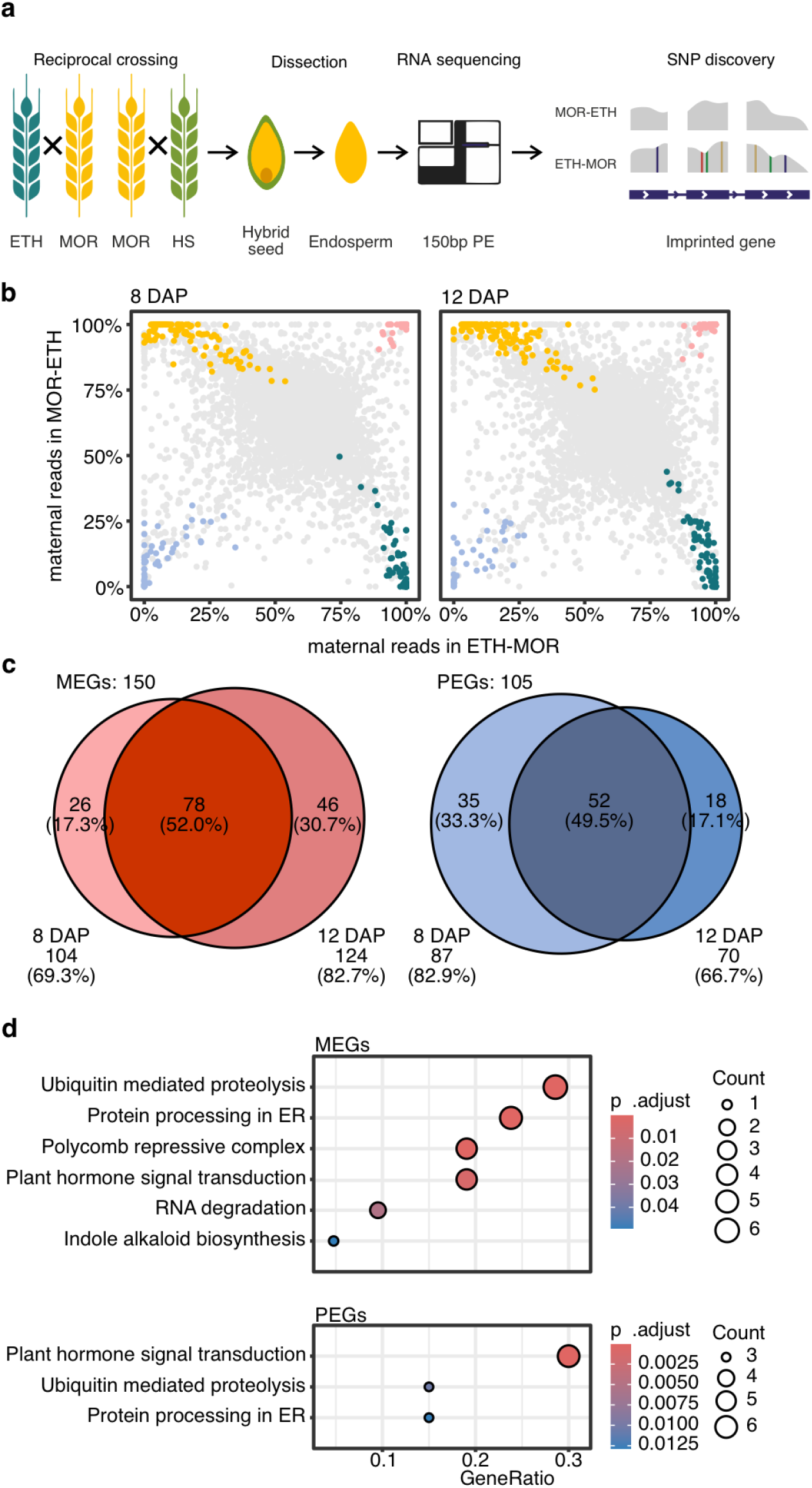
Identification of 255 imprinted genes in barley endosperm. (**a**) Experimental workflow. (**b**) Visualization of the candidate-imprinted and genotype-specific genes in F_1_ MOR-ETH reciprocal crosses 8 and 12 days after pollination (DAP). Dots: pink/blue – maternally/paternally expressed genes (MEGs/PEGs, respectively); yellow/dark green – genes with genotype (MOR/ETH)-specific expression, respectively; grey – non-imprinted genes with biallelic expression. (**c**) Venn diagrams of the MEGs (red) and PEGs (blue) 8 and 12 DAP. (**d**) Bubble plot of KEGG pathway enrichment among MEGs and PEGs. The color scales indicate FDR-adjusted p-values, and the sizes of bubbles indicate gene counts.

In terms of functions associated with imprinted genes (KEGG database; Fig. 1d, Supplementary Data Set 6), MEGs were enriched in ubiquitin-mediated proteolysis via the Skp1-Cullin-F-box complex (Supplementary Fig. 4a). Both MEGs and PEGs were enriched in plant hormone signal transduction, with MEGs involved in auxin and abscisic acid (ABA)-based regulation, and PEGs in ABA and cytokinin signaling (Supplementary Fig. 4b). Cytokinin signaling-associated PEGs included four authentic *HISTIDINE-CONTAINING PHOSPHOTRANSFER PROTEINs* and a *B-type TWO-COMPONENT RESPONSE REGULATOR* transcription factor. This points to parent-of-origin-specific effects in hormonal regulation and proteolysis during mid-to-late stages of barley endosperm development.

To assess the degree of evolutionary conservation, we identified orthogroups for imprinted genes of barley, maize, rice and wheat^7–9,14,15^. A total of 42 orthogroups with 120 genes were conserved in two species, and 2 orthogroups (8 genes) in three species (Fig. 2a, Supplementary Data Set 7). Surprisingly, only 31 imprinted barley genes (12.2%) were orthologous to imprinted genes in any other cereal and only 17 to hexaploid wheat (*Triticum aestivum*). Comparison of barley with the wheat progenitors, *Aegilops tauschii* and *Triticum turgidum*, revealed 21 and 22 conserved imprinted genes, respectively (Fig. 2b, Supplementary Data Set 8). However, 14 and 15 imprinted genes shared between barley and *A. tauschii* or *T. turgidum*, respectively, were not conserved between barley and *T. aestivum*. Among the 31 conserved imprinted barley genes, 14 genes, including DOF, MYB, and NAC transcription factors, were expressed in endosperm but silent in embryo and maternal tissues (MTs), while the rest were widely expressed in embryo and/or grain maternal tissues (Fig. 2c, Supplementary Data Set 9). These data suggest a low degree of conservation of the imprinted genes in cereals, thus supporting data from dicots^16^. We hypothesized that the parent-of-origin-specific effects might be better conserved at the pathway level. In total, 121 biological processes (BPs), 73 molecular functions (MFs) and 35 cellular components (CCs) were common for the imprinted genes in maize, rice, wheat, and barley (Fig. 2d, Supplementary Data Set 10). Exclusion of general and narrow terms revealed a core conserved set of 8 BPs, 4 MFs and 1 CC (Fig. 2e). The hormonal phosphorelay signal transduction system was imprinted across cereals and significantly enriched in barley (FDR-adjusted p < 0.001, 6.85-fold enrichment), with 10 genes being specifically involved in the two-component system (TCS) pathway of cytokinin signaling (Supplementary Data Set 10). Activity of the TCS genes was previously localized to barley endosperm transfer cells (ETCs), suggesting an important role for imprinting in endosperm development and grain filling^17^.

**Fig. 2.**
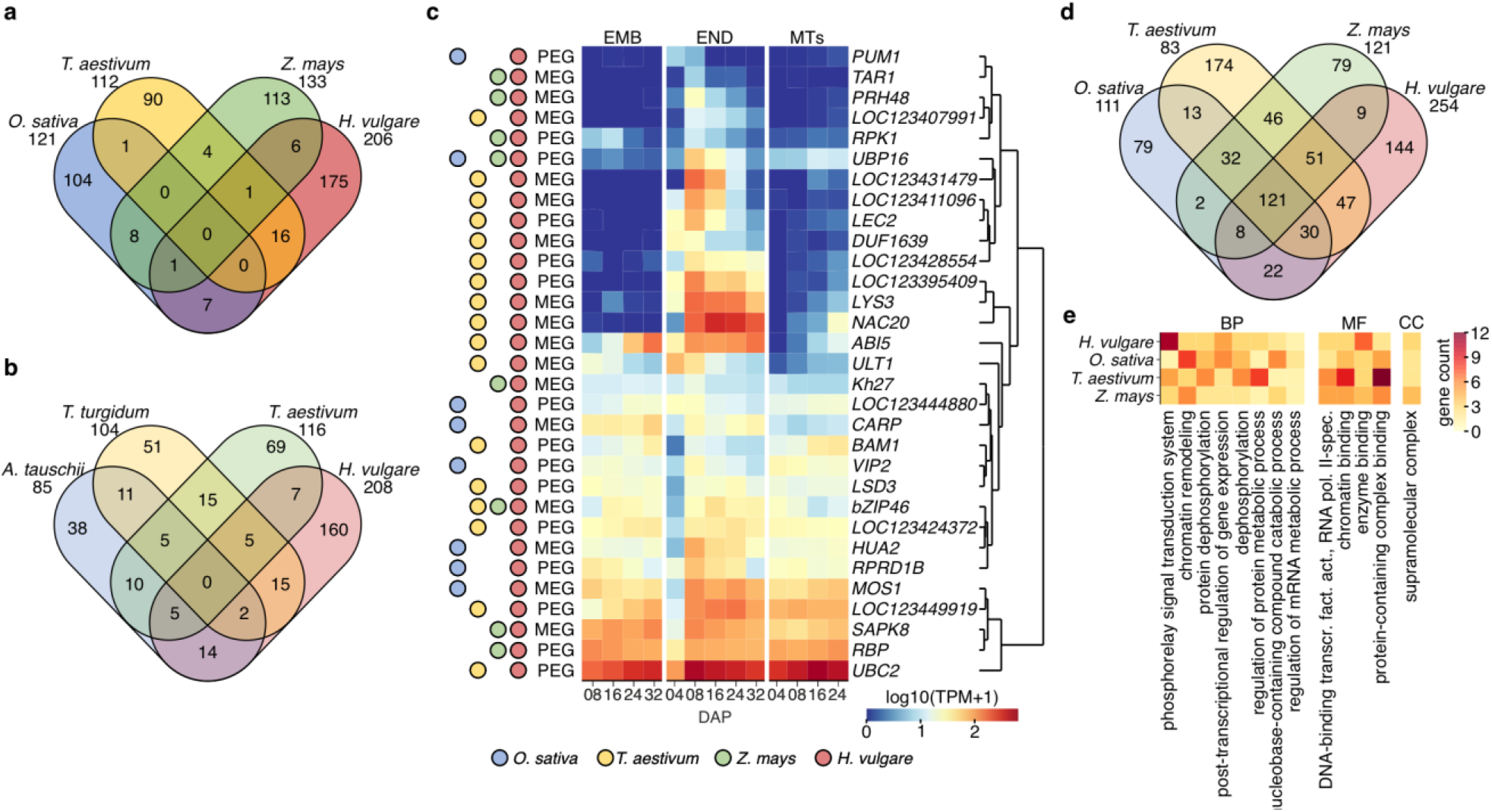
Genomic imprinting operates via conserved biological processes in cereals. (**a**) Venn diagram of imprinted orthogroups in cereals. Non-barley data are from^7–9,14,15^. (**b**) Venn diagram comparing imprinted genes found in *Triticeae* species. Non-barley data are from^9^. (**c**) Expression-based hierarchical clustering of barley maternally and paternally expressed genes (MEGs and PEGs, respectively) found imprinted in more than one species. The total expression is shown at different days after pollination (DAP) in barley endosperm (END), embryo (EMB) and maternal tissues (MTs)^18^. Genes identified as imprinted in different cereals are indicated by the color circles on the left. (**d**) Gene ontology-based biological processes represented by imprinted genes shown in (a). (**e**) Numbers of imprinted genes per gene ontology-based biological process (BP), molecular function (MF) and cellular component (CC) found in all four cereal species filtered for term size and similarity.

To explore the temporal expression pattern of imprinted genes, we performed k-means clustering of them using an endosperm transcriptomic data series from 4 to 32 DAP^18^ and identified three co-expression clusters (CL). The 56 CL1 genes (36 MEGs, 20 PEGs) and 97 CL2 genes (53 MEGs, 44 PEGs) peaked at 4 and 8 DAP, corresponding to the coenocytic and cellularizing stages, respectively (Fig. 3a, Supplementary Data Set 5). This indicated that although our samples represented cellularizing and cellularized endosperm, they contained also some imprinted genes peaking in the coenocytic endosperm. The expression profile of 102 CL3 genes (61 MEGs, 41 PEGs) increased from 4 to 8 DAP and remained high until 32 DAP. These genes may remain imprinted beyond 12 DAP (as reported for some genes in wheat) or may transition to biased or biallelic expression (as found in maize)^7–9,14,15^.

**Fig. 3.**
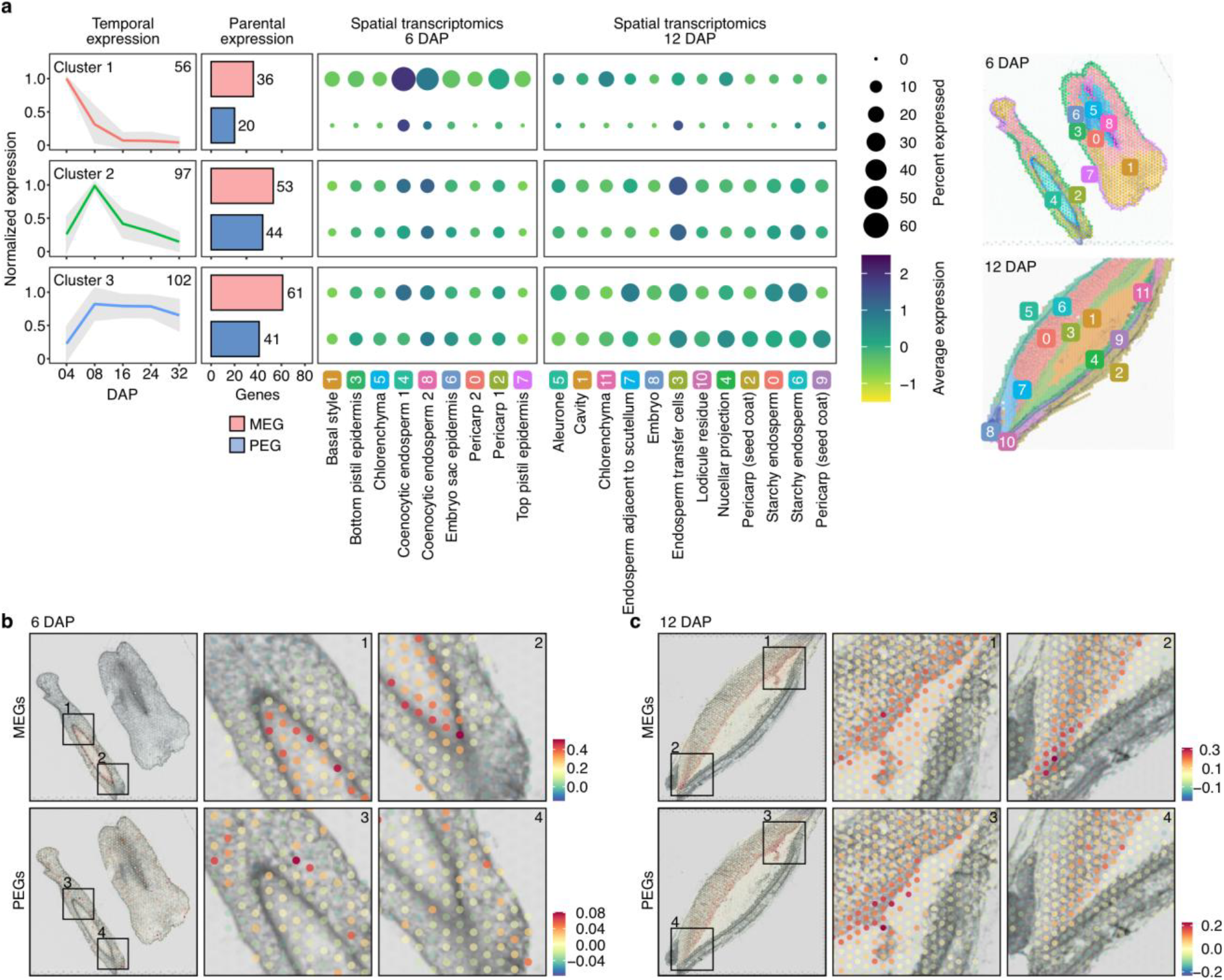
Spatio-temporal dynamics of expression of barley imprinted genes. (**a**) K-means-based temporal expression clusters of 255 barley imprinted genes. Number of genes per cluster is given in the top right corner. Mean expression (colored line) and standard deviation (grey area) in 4 to 32 days after pollination (DAP) endosperm are based on^18^. Parental expression graphs give the numbers of maternally/paternally expressed genes (MEGs/PEGs) per k-means cluster. Spatial transcriptomics plots indicate average expression of imprinted genes in 6 and 12 DAP grain sections. Circle sizes - percentage of expression; color - z-score expression; upper row – MEGs; lower row - PEGs. The x-axis represents individual tissues as indicated in the representative sections on the far-right based on^20^. (**b-c**) Spatial cumulative expression of 70/84 MEGs, and 69/71 PEGs at (b) 6 and (c) 12 DAP, respectively. Color scales represent the normalized counts with the cut-offs individually specified on the image. Insets show detailed views of the spatio-temporal hotspots.

The expression of imprinted genes in Arabidopsis coenocytic endosperm is concentrated at the chalazal pole^19^. The absence of a chalaza in cereals let us also analyze the expression of imprinted gene clusters in a spatially-resolved dataset of developing barley grains^20^ (Fig. 3a). The expression of CL1 genes localized mainly to the periphery of the coenocytic endosperm at 6 DAP (Fig. 3b; Supplementary Fig. 5). The most expressed CL1 genes were 5 members of the *non-specific LIPID TRANSFER PROTEINS* (*nsLTPs*) involved in cuticle formation^21^. Interestingly, CL2 genes were highest in ETCs at 12 DAP and included ABA and cytokinin signaling and regulatory genes (Fig. 3a and c, Supplementary Fig. 6). ETCs represent nutrient-absorbing tissue directly adjacent to the nucellar projection, a tissue of maternal origin with a key role in transporting nutrients to developing endosperm^22^. The analogous functions of the chalazal endosperm and ETCs suggest convergent evolution of expression of imprinted genes at the interface between the maternal and the filial bodies. Such patterning may reflect parent-of-origin-specific competition for maternal resources and supports the parental conflict hypothesis^23^.

The expression of CL3 genes was scattered across the entire endosperm (Fig. 3a, Supplementary Fig. 7). CL3 included the prolamin box binding transcription factor *HIGH LYSINE 3* (*LYS3*; LOC123395703), known to positively regulate hundreds of genes involved in endosperm filling and maturation^24^. Grains of its loss-of-function mutants have shrunken endosperm, reduced starch and HORDEINs, and higher lysine content^25^. However, *lys3* phenotypes segregate in a 3:1 recessive Mendelian manner^25,26^, thus deviating from 1:1 or 2:1:1 ratio typical for imprinted genes^27^. To explain this discrepancy, we analyzed heritability of *lys3* phenotypes over grain development using molecular and histological methods. Hierarchical clustering of genome-wide endosperm transcriptomic samples from all F_1_ hybrid combinations of wild-type (L) and *lys3*.*c* (l; E37*) plants revealed a 1:1 segregation at 12 DAP, separating the samples with maternally-inherited *LYS3* wild-type (LL-L and LL-l) and mutant (ll-L and ll-l) endosperm constitutions (Fig. 4a, Supplementary Data Set 11). Interestingly, the 20 DAP segregation pattern corresponded to 3:1, with separated homozygous mutant samples (Fig. 4b). The maternal allele effects were obvious also when comparing the number of genes downregulated in crosses with maternal wild-type (LL-l) or maternal mutant (Il-L and ll-l) endosperm constitution relative to homozygous wild-type (LL-L) at 12 DAP (Fig. 4c). By comparing genes downregulated in ll-L and ll-l relative to LL-L endosperm over time, we identified 90 *LYS3* targets downregulated in 12 DAP ll-L (Fig. 4d; Supplementary Fig. 8a, b). These genes showed up to 5.6-fold enrichment for the GO terms *endopeptidase regulator activity* and a striking 40-fold enrichment for *nutrient reservoir activity* (Fig. 4e, Supplementary Data Set 12). The *endopeptidase regulator activity* included genes coding for various enzyme inhibitors and SERPINs Z4/Z7, representing the major components of endosperm albumin with storage and serine protease inhibitory functions^28,29^. The *nutrient reservoir activity* genes coded for the major storage proteins HORDEINs and GLUTENINs^30,31^, showing 1:3 expression pattern at 8 DAP, 1:1 maternal allele-dependent pattern at 12 DAP, and 3:1 homozygous mutant-dependent pattern (particularly for *B1-hordein-like* genes) at 20 DAP (Fig. 4f, Supplementary Data Set 13). Therefore, *LYS3* temporally regulates its target genes in a non-Mendelian, maternal allele-dependent manner.

**Fig. 4.**
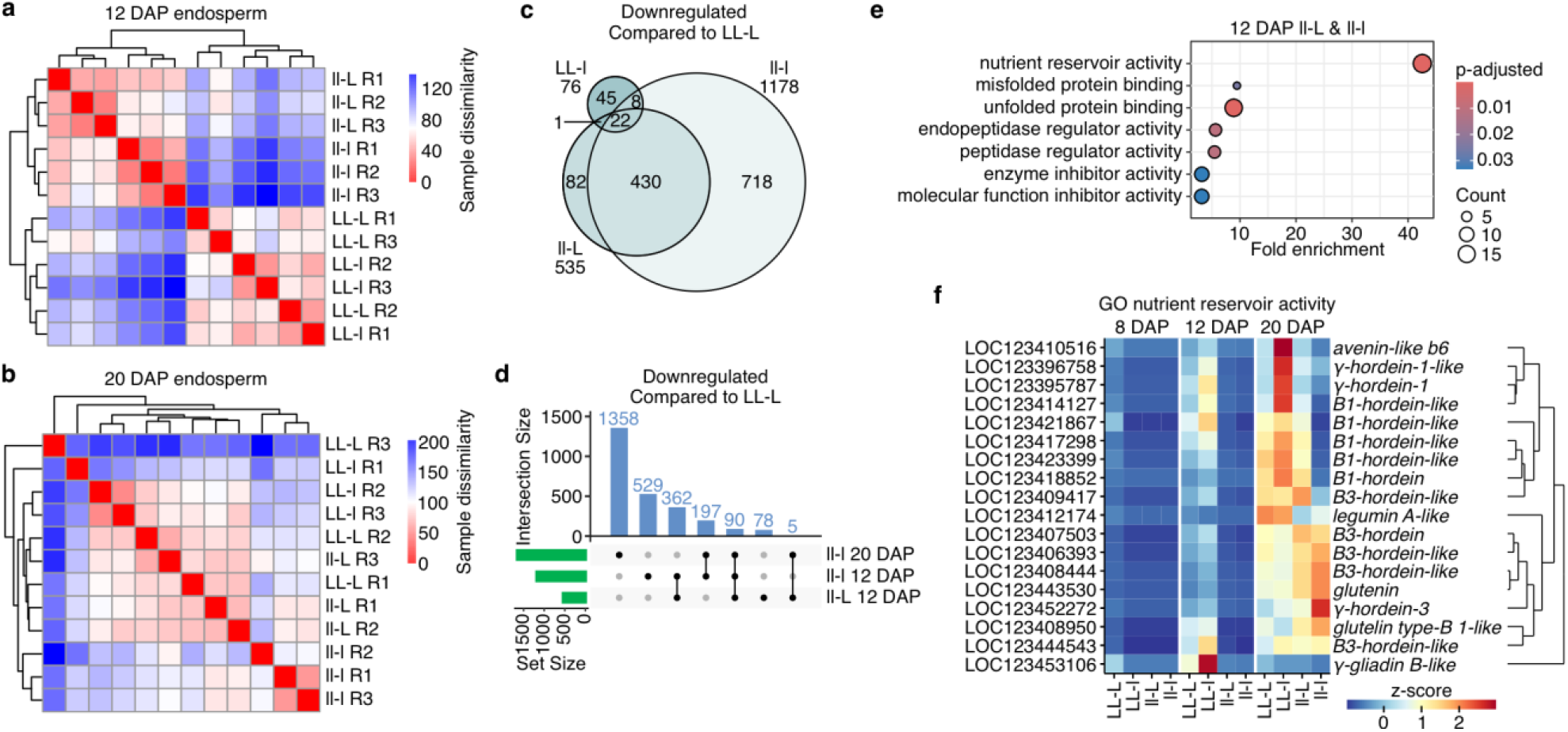
Non-Mendelian effects of *LYS3* on endosperm transcriptome. (**a-b**) Hierarchically clustered sample dissimilarity heatmaps of endosperm transcriptomes of the *LYS3* (L) and/or *lys3*.*c* (l) crosses at (a) 12 and (b) 20 days after pollination (DAP). Maternal alleles are given first. R – biological replicates. (**c**) Venn diagram of significantly downregulated (FDR-adjusted p < 0.05, log2FoldChange < −1) genes in individual crosses at 12 DAP compared to LL-L endosperm. (**d**) UpSet plot of significantly downregulated genes in ll-L and ll-l crosses at 12 and 20 DAP. The x-axis set size defines a total number of differentially expressed genes (DEGs) per cross. The y-axis shows the number of DEGs (intersection size) unique (single dots) or shared (connected dots) among crosses. (**e**) Bubble plot of GO-term enrichment of common significantly downregulated genes in ll-L and ll-l crosses relative to LL-L at 12 DAP. The x-axis shows fold-enrichment, the color scale indicates FDR-adjusted p-value, and size of the bubbles indicate gene count. (**f**) Hierarchically clustered expression of GO term nutrient reservoir activity genes from (e) in different cross combinations at 8, 12 and 20 DAP. Color scale indicates z-score.

Since LYS3 regulates both storage protein and starch synthesis^32^, we analyzed the genetic basis of their accumulation during grain filling. HORDEINS were abundant in LL-L and LL-l and significantly reduced in ll-L and ll-l endosperms at 16 DAP (two-way ANOVA with Tukey’s HSD, p < 0.05), resembling a 1:1 segregation (Fig. 5a-b, Supplementary Fig. 9). At 20 DAP, the ll-L cross endosperm resembled the wild-type sample, suggesting a shift towards a 3:1 ratio. Starch accumulated from 12 DAP with similar numbers of starch granules per endosperm, but with different granule morphology (Fig. 5c-d, Supplementary Fig. 10, Supplementary Data Sets 14 and 15). Starch granules of LL-L and LL-l crosses were larger and more uniform in size and shape, while the ll-L and ll-l combinations contained smaller and less uniform granules, corresponding to maternal allele-dependent 1:1 segregation. The segregation pattern changed to 3:1 at 20 DAP, with more uniform and lens-shaped starch granules in all combinations except ll-l. Interestingly, the number of starch granules per endosperm correlated negatively with the number of *LYS3* alleles (R^2^ = 0.9, p < 0.001) at 20 DAP, suggesting a quantitative effect of *LYS3* expression (Fig. 5d). Finally, we performed high-throughput quantitative analysis of the shrunken endosperm phenotype by measuring and scoring dry grain thickness profiles of self-pollinated wild-type, homozygous *lys3*.*c*, and self-pollinated heterozygous *LYS3*/*lys3*.*c* plants (Fig. 5e, Supplementary Fig. 11a-b, Supplementary Data Sets 16 and 17). A Gaussian mixture model fitted to the observed score distribution of the heterozygote-delivered F_2_ seeds strongly preferred two components over three (ΔBIC = 19.0), indicating no detectable intermediate phenotypic class and excluding semi-dominant 2:1:1 segregation (Supplementary Fig. 11c). Simulating 1000 synthetic F_2_ datasets under 3:1 segregation and comparing them to the observed distribution confirmed that the observed F_2_ score distribution matches the 3:1 expectation (Kolmogorov-Smirnov test, p = 0.165; Supplementary Fig. 11d).

**Fig. 5.**
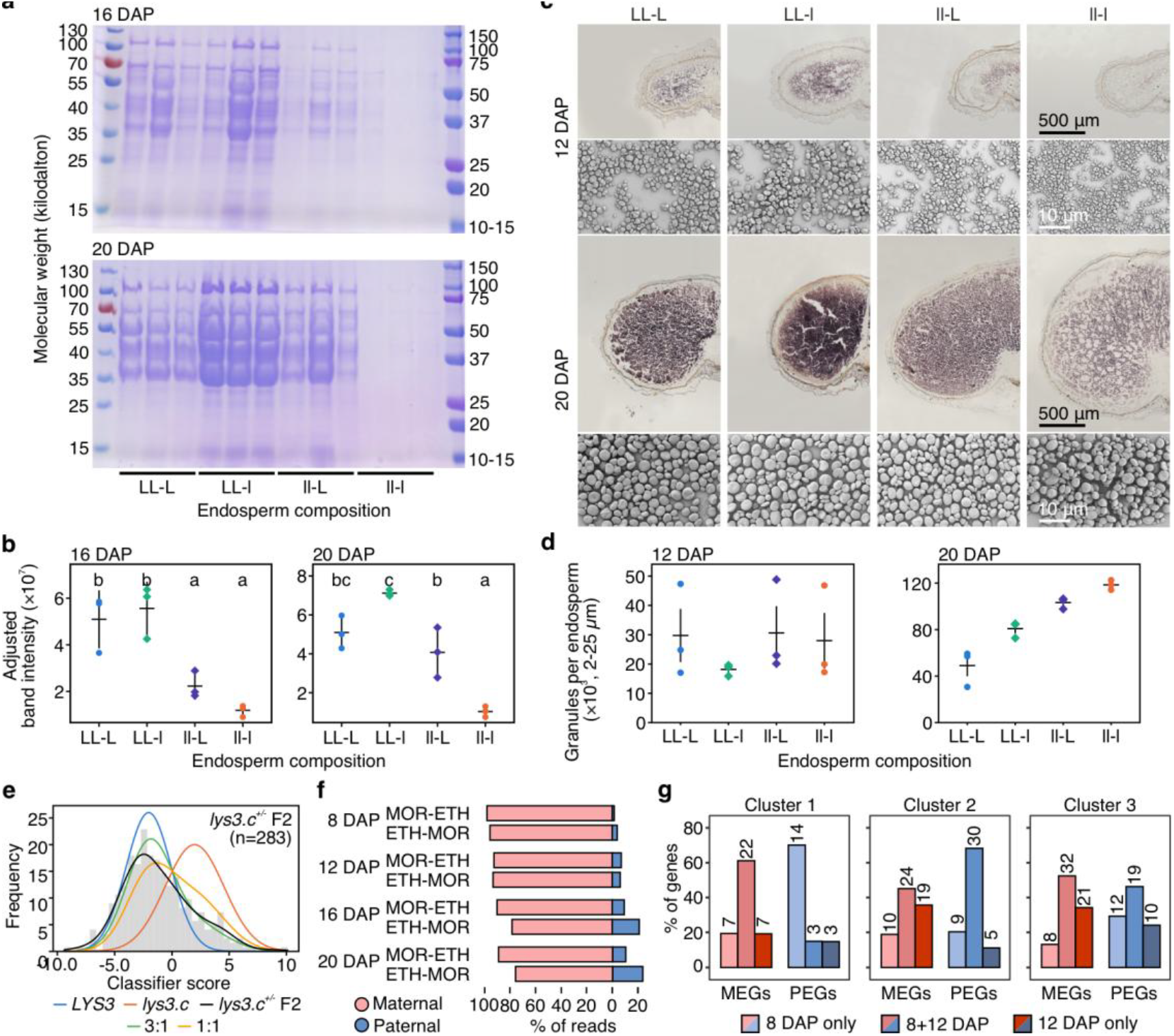
Maternal *LYS3* allele controls accumulation of storage proteins and starch. (**a**) SDS-PAGE of HORDEINs isolated from grains of *LYS3* (L) and *lys3*.*c* (l) crosses at given days after pollination (DAP). Equivalent fractions of total extracted proteins were loaded. (**b**) Difference in band intensities in individual crosses. Values marked with the same letter do not differ within the respective group according to two-way ANOVA with Tukey’s HSD (p < 0.05). (**c**) Accumulation of starch granules (brown) based on Lugol staining of transverse grain sections of individual crosses at given DAPs, and morphology of starch granules obtained by scanning electron microscopy at 12 and 20 DAP. (**d**) Quantification of starch granules per endosperm of individual *LYS3* and *lys3*.*c* crosses at given DAPs. (**e**) Machine learning classification of thickness profiles of individual F_2_ grains from self-pollinated *lys3*.*c* heterozygotes. Blue and red line indicate peak positions of homozygous *LYS3* and *lys3*.*c* genotypes from training dataset. Green and yellow dashed line correspond to distribution of synthetic F_2_ datasets under 3:1 and 1:1 segregation, respectively. (**f**) Percentage of maternal (salmon) and paternal (blue) *LYS3* transcripts based on RT-qPCR from reciprocal F_1_ MOR-ETH crosses at given DAPs. (**g**) Time specificity of imprinted genes from expression clusters 1 to 3 in 8 and 12 DAP RNA-seq data.

To explain the shift in expression patterns of LYS3 targets, we analyzed expression of the parental *LYS3* alleles. Based on RNA-seq of F_1_ MOR-ETH and MOR-HS endosperms, only the maternal *LYS3* was expressed at 8 DAP, but the paternal allele became partially active (up to 18%, compared to 33.3% expected reads) at 12 DAP (Supplementary Fig. 12, Supplementary Data Sets 1-4). This was confirmed on an extended timescale (8, 12, 16 and 20 DAP) by reverse transcription quantitative PCR (Fig. 5f), which revealed an increasing amount of paternal *LYS3* transcript over time, reaching 10-23% of the amplicons. Hence, the initial maternal expression of *LYS3* at 8 DAP was followed by partial transcriptional activation of the paternal allele at later stages of endosperm development, consistent with a progressive compensation for the early maternal allele deficiency and the subsequent transition toward a recessive Mendelian segregation frequency. Taken together, the 12-16 DAP transcriptomes of *LYS3*/*lys3* crosses, their HORDEIN phenotypes, and the dynamics of paternal *LYS3* allele expression explain why *lys3* shows an early maternal 1:1 effect during grain filling but a recessive 3:1 phenotype in mature F2 grains.

Asking whether this changing expression pattern of paternal genes is common in barley, we analyzed parent-of-origin specific expression of 102 genes from CL3 (Fig. 5g, Supplementary Data Set 5). In total, 52.5% MEGs and 46.3% PEGs retained their character at both 8 and 12 DAP (Fig. 5g). On the contrary, 13.1% MEGs and 29.3% of PEGs were parent-of-origin-expressed at 8 DAP but parent-of-origin biased or biallelically expressed at 12 DAP. The opposite case of biparental expression at 8 DAP and parent-of-origin specific expression at 12 DAP was represented by 34.4% MEGs and 24.4% PEGs. The latter case suggested that profiling at late stages of endosperm development may reveal novel imprinted genes possibly affecting the final stages of grain maturation. Indeed, 83 imprinted genes were detected in fully developed and desiccating hexaploid bread wheat endosperm at 25 DAP^9^.

Our combined transcriptome and genetic data show that genomic imprinting, previously well documented during early endosperm development across different species, has also an important role during grain filling and maturation in barley. The transition from initial maternal expression of transcription factor gene *LYS3*, regulating nutrient and energy supply in multiple ways, to later activation of the paternal allele, suggests a tight maternal genome-exerted control over the developing grain, in line with the parental conflict theory^23^.

### Online content

Any methods, additional references, Nature Portfolio reporting summaries, source data, extended data, supplementary information, acknowledgements, peer review information; details of author contributions and competing interests; and statements of data and code availability.

## Data availability

All data supporting this study are included in the article and its Supplemental material. Sequence data from this article can be found in the Gene Expression Omnibus under accession numbers GSE349961 and GSE350207.

## Code availability

The central code repository for the GATK output processing and post-processing analyses is available via GitHub at https://github.com/pecinkagroup/barley-imprinting/. Post-processing analyses were implemented with basic R (v.4.5.2) functionality using publicly available R packages.

## Acknowledgments

This work was supported by the Ministry of Education Youth and Sports of the Czech Republic grants LUC24011 and “TowArds Next GENeration Crops (TANGENC)” (No. CZ.02.01.01/00/22_008/0004581) and MSCA Doctoral Network HeatDDR (Funded by the European Union under Grant Agreement No. 101169110). Computational resources were supplied by the project “e-Infrastruktura CZ” (e-INFRA CZ LM2018140) supported by the Ministry of Education, Youth and Sports of the Czech Republic. Barley FIND-IT screening and variant isolation was performed with funding from the Carlsberg Foundation (grant CF15-0236). We thank H. Tvardíková, K. Lakomčíková and M. Krautsova for assistance, Z. Bursová for plant care, P. Navrátil for IT support, JIC Bioimaging facility and staff for their contribution to this publication and O. Mittelsten Scheid for comments.

## Author contributions

A.P., M.K. and C.L.P conceived and designed the project. A.P. supervised the project, M.K., A.L., A.N., P.Š., Z.K., K.K., K.T., B.S., T.D., M.Č. performed experiments, M.P.-L. and J.W. provided spatial transcriptomics data; M.W.R, S.K., C.D. provided the barley FIND-IT variant; A.P and M.K. wrote the manuscript with contribution of all authors. All authors have seen and agreed upon the final version of the manuscript.

## Additional information

### Supplementary information

The online version contains supplementary material available at

