## Supplementary material for "Temporal genomic imprinting regulates grain filling in barley": Materials and Methods

Experiments within this study did not require ethics board approval.

### Plant materials and growth conditions

Barley (*Hordeum vulgare* L. subsp. *vulgare*) genotypes Morex (MOR; BCC906), Ethiopian landrace (ETH; HOR10350), Bomi (NGB5096), *lys3.c*$^{E37*}$ (also known as Risø19) mutant in Bomi background[1], RGT Planet, *lys3*$^{W70*}$ mutant in RGT Planet background[2] (*lys3.e*; CB-FINDit-Hv-048) and wild barley *H. vulgare* subsp. *spontaneum* (K. Koch) Asch. & Graebn. (HS; HOR12560), were used throughout this study. The CB-FINDit-Hv-048 *lys3.e* line was obtained from the previously described RGT Planet FIND-IT mutant collection, the allele was identified by ddPCR screening using Bio-Rad Unique Assay ID dMDS356047506[2]. The *lys3.c* and *lys3.e* plants were genotyped for the mutations using dCAPS markers and validated using Sanger sequencing (Supplementary Table 1).

For plant growth, the grains were spread on a wet cellulose tissue paper covered with one layer of filter paper, stratified at 4 °C in the dark for 48 h, and germinated in the dark at 25 °C for 3 days. Two grains per genotype were germinated weekly for six weeks to obtain plants of different genotypes flowering at the same time. Germinating grains were planted into 5 × 5 cm peat pots containing a mixture of soil and sand (2:1; v/v) and were grown in a phyto chamber under a long-day regime (16 h day 15 °C; light intensity 200 μmol $m^{-2}$ $s^{-1}$; 8 h night 13 °C, humidity 60%). After two weeks, plants were transferred to 15 × 15 cm pots containing the same soil and cultivated until flowering. Two-week-old HS plants were vernalized under a short-day regime (8 h day 4 °C; light intensity 200 μmol $m^{-2}$ $s^{-1}$; 16 h night 4 °C, humidity 85%) for three weeks. Vernalized HS plants were transferred to 15 × 15 cm pots containing the same soil and cultivated in the long-day conditions until flowering.

Emasculation was performed by the Bonnett method[3]. Individual spikes were packed in glassine bags and labeled. Pollination was performed at the W10 stage[4], i.e. typically 2 to 5 days after emasculation. Mature anthers from the pollen donor were introduced through the incision in the lemma onto the stigma of the mother plant using fine forceps.

The F1 hybrid grains were collected at the desired DAP. Endosperm was manually dissected and three to five endosperms were pooled per biological replicate. Each sample was checked for purity by flow-cytometric ploidy measurement as described[5]. The samples were flash frozen in liquid nitrogen and stored at −80 °C until use.

**RNA extraction and sequencing**

Total RNA was isolated using Monarch® Total RNA Miniprep Kits (New England Biolabs) according to the manufacturer's instructions, including on-column DNase I (Sigma-Aldrich) treatment. RNA quality was monitored using a Bioanalyzer 2100 with RNA 6000 Pico Chips (all Agilent). Samples with RNA integrity number >6.8 were processed into RNA-seq libraries using NEBNext® Single Cell/Low Input RNA Library Prep Kit for Illumina® with poly-A selection. The mRNA-enriched libraries were sequenced as paired-end 150 bp RNA-seq reads on a NovaSeq 6000 instrument (Illumina).

**Variant calling and identification of uniparentally-expressed genes**

Transcriptomic data were processed using standard GATK pipeline with minor modifications (Supplementary Figure 13). Raw reads were trimmed for adaptors by flexbar[6] and aligned to the Morex reference genome v3[7] using BWA version 0.7.17[8] with a smart pairing option for paired end alignment. Duplicates were removed and SNPs were called using GATK version 4.2.5.0[9]. The obtained SNPs were filtered by genotype quality (GQ > 19). NCBI *Hordeum vulgare* subsp. *vulgare* Refseq Annotation Release 100 was used to assign individual SNPs to annotated features[10]. Exonic SNPs with only one alternative allele were used to determine expression patterns. Each SNP was tested for significant deviation (p-adj < 0.05) from the expected 2:1 ratio of the maternal to paternal allele-derived reads in the reciprocal crosses using Binomial test. The gene was considered as candidate MEG or PEG based on at least one exonic SNP with the same and significant bias in the reciprocal cross combinations, and the proportions of sequencing reads differed at least five-times between the maternal and paternal alleles in both direction of the reciprocal crosses. In case of multiple diagnostic SNPs, we required the bias to be consistent for all of them. Differential expression analysis was further employed to filter out candidate endosperm MEGs and PEGs for potential tissue cross-contamination using published grain transcriptome dataset GSE233316[11]. Using DESeq2 package v.1.38.3[12] in R v.4.2.2 software[13] candidate imprinted genes were tested for significant upregulation at 8 DAP endosperm. Candidate MEGs were tested in reference to 8-16 DAP embryo and 4-16 DAP maternal tissues, candidate PEGs were tested in reference to 8-16 DAP embryo (Benjamini-Hochberg FDR-adjusted p-value <0.05). Venn diagrams were drawn using the eulerr v.6.1.0 package in R[14].

**Validation of allele-specific expression**

Independent hybrid endosperm samples were prepared (using protocol described above) to validate parent-of-origin-specific expression of selected candidate genes. Total RNA was reverse transcribed into cDNA using RevertAid H Minus First Strand cDNA Synthesis Kit (Thermo Fisher Scientific). Primers were designed to amplify 200–1,100 bp fragments of the candidate genes harboring informative SNP(s) (Supplementary Table 1) using a standard PCR. The amplicons were gel purified using GeneJet Gel Extraction Kit (Thermo Fisher Scientific) and subjected to Sanger sequencing followed by *in silico* analysis using SnapGene v6.2 software (GSL Biotech LLC).

**Gene ontology and KEGG pathway enrichment**

GO terms provided in NCBI *Hordeum vulgare* subsp. *vulgare* Annotation Release 100 were used to generate a custom-made GMT file and enrichment analysis was performed by the g:Profiler online tool[15]. KEGG pathway enrichment was performed using clusterProfiler version 4.16.0[16]. KEGG annotation for NCBI gene models was obtained using KEGGREST version 1.50.0 packages in R[16,17]. The list of pathways was filtered for Benjamini-Hochberg adjusted p-value < 0.05 and visualized using dotplot in R.

**Evolutionary conservation**

GeneIDs of imprinted genes identified in *Oryza sativa*[18,19], *Triticum aestivum*[20] and *Zea mays*[21,22] were converted to recent annotations using ID History Converter Tool at Ensembl Plants Release 62[23]. Groups of orthologous genes (orthogroups) were identified using protein sequences from *Hordeum vulgare* subsp. *vulgare* Refseq Annotation[10], *O. sativa* IRGSP v1.0 annotation[24], *T. aestivum* IWGSC Refseq Annotation v1.1[25] and *Z. mays* B73 RefGen v5 Annotation[26]. Orthogroups were assigned to individual imprinted genes and compared across different species. For functional conservation, assigned GO terms for Biological processes (BP), Molecular functions (MF) and Cellular components (CC) were retrieved for individual genes using g:Profiler[15], and reduced using REVIGO[27], both with default parameters. Number of genes for individual GO terms were collected per species. To get the most meaningful terms, the following filters were applied: term size more than or equal to 10 and less than or equal to 500, the number of species with a gene with a specific GO term is equal to 4, and semantic similarity threshold (dispensability) in all species is less than or equal to 0.7.

**Analysis of spatio-temporal expression**

To get temporal information about imprinted gene expression, MEGs and PEGs were clustered based on their expression patterns in developing barley endosperm[11] using the Hartigan-Wong k-means clustering algorithm with 1,000 iterations in R[13]. To investigate an optimal number of clusters, the total sum of squares within clusters was inspected. Spatial transcriptomics data of developing barley grain[28] were obtained and individual imprinted genes detected in the dataset were combined into single SpatialFeaturePlot per MEGs and PEGs respectively, using Seurat v5.4.0[29]. Average and Percent expression in individual grain compartments were plotted for individual genes using the provided tissue annotation, and mean of Average and Percent expression in individual grain compartments were plotted for MEGs and PEGs per kmeans cluster.

**Transcriptome analysis of *lys3.c* crosses**

The raw reads were trimmed for adaptors by Trim Galore v.0.4.1[30] and aligned to the *H. vulgare* cv. Morex reference genome v3[7] using HiSat2 v.2.1.0 genomic mapper[31]. Aligned reads were assigned to features and meta-features using Subread v.1.5.2 software[32] according to the NCBI *Hordeum vulgare* subsp. *vulgare* Annotation Release 100. Differential expression analysis was performed using the DESeq2 v.1.24.0 package[12] in R v.3.6.3 software[13]. A gene was declared to be significantly differentially expressed if the Benjamini-Hochberg FDR-adjusted p-value was <0.05. The PCAs were done after applying variance stabilization transformation[33]. Venn diagrams were drawn using the eulerr v.6.1.0 package in R[14].

**Allele-specific reverse transcription quantitative PCR**

Allele-specific expression levels were determined using reverse transcription-qPCR (RT–qPCR). Allele-specific primers were designed to include cultivar-specific SNPs, and the specificity of primers was tested using qPCR on DNA from individual cultivars. RT–qPCR was performed using qPCR 5x HOT FIREPol® EvaGreen® qPCR Mix Plus (Solis BioDyne OÜ) on the CFX96TM Real-Time PCR Detection System (Bio–Rad). The data were analyzed using the $2^{-\Delta\Delta Cq}$ method with CFX Maestro 2.0 software (Bio–Rad). Three biological replicate PCR amplifications were performed for each sample. The expression level was standardized against the reference *E3 UBIQUITIN LIGASE* with a stable expression between 8 and 20 DAP. Sequences of all primers used for RT-qPCR are listed in Supplementary Table 1.

**SDS-PAGE**

HORDEINs were extracted with minor modifications to a previously described protocol[34]. Whole grains 12, 16, or 20 DAP were flash-frozen in liquid nitrogen and ground using a mortar and pestle. Approximately 40 mg of fine powder was extracted in 1 mL of 60% (v/v) isopropanol and 2% (v/v) β-mercaptoethanol in 50 mM Tris-Borate buffer (pH 8), by heating at 60 °C for 1 h (1500 rpm).

After centrifugation (17,000 *g*, 10 min), two 0.525 mL aliquots were separately mixed with 20% (v/v) NaCl solution and precipitated overnight at -20 °C, centrifuged (17,000 *g*, 2 min) and supernatant was discarded. Pellets were air-dried and dissolved in 50 µL Sigma Total Plant Protein Reagent Type 4 containing 1% cOmplete™, EDTA-free Protease Inhibitor Cocktail. Proteins were denatured by mixing samples in volume ratio 4:1 with 4× Laemmli Sample Buffer containing β-mercaptoethanol (in a 9:1 ratio) at 70 °C for 10 min. An equal volume of each sample was loaded onto 12 % SDS-PAGE gel (TGX™ FastCast™ Acrylamide Kit, BioRad) using Dual Color Standards (Bio-Rad) and PageRuler™ Plus Prestained Protein Ladder (Thermo Scientific). Developed gels were fixed in 50% methanol/10 % acetic acid solution for 15 min. Gels were stained in Coomassie brilliant blue solution for 1h and destained (10 % methanol, 10 % acetic acid) four times, each for 20 min, and left in water overnight. Gel images were captured and signals quantified using Amersham Imager 600 (Cytiva) or Azure 600 (Azure biosystems).

**Grain sectioning and starch staining**

Grains were harvested 8, 12, 16 and 20 DAP in at least three biological replicates, fixed with a freshly prepared 4% (w/v) paraformaldehyde, with 2% (v/v) Tween-20 and 2% (v/v) Triton X-100 (pH 7, adjusted by HCl). To increase fixation efficiency, the vacuum was broken every 10 min and then reapplied for a total of six cycles. Subsequently, grains were transferred into new fixative (as described above), stored overnight at 4°C, dehydrated using ethanol series, cleared by ROTIHistol (Carl ROTH) series, and embedded into Paraplast (Sigma-Aldrich). Transverse sections of 10 µm were cut with a Reichert-Jung 2030 microtome and attached to Adhesion Slides Superfrost Ultra Plus (Thermo Fisher Scientific).

The sections were pretreated using ROTIHistol, rehydrated by ethanol series and post fixed in 4% (w/v) paraformaldehyde. Dehydrated sections were stained in 1% (w/v) iodine and 1% (w/v) potassium iodide for 30 s, rinsed in sterile water and embedded in Entellan (Sigma-Aldrich). Images were taken with a light microscope (Olympus BX60) equipped with a Promicra 3-3CP camera (Sony). The grains were captured in segments using QuickPHOTO CAMERA 4 (Promicra). To generate complete grain images, the segments were merged using Fiji[35].

**Quantitative starch extraction**

Endosperms were dissected from developing grains collected at 12 and 20 DAP and stored at -80°C. For each sample, 3 to 5 individual grains were thawed on ice in a 1.5-ml tube prior to dissection. After homogenization using a plastic pestle in 300 µl ice-cold water, the homogenate was centrifuged at 10,000 g, 4°C for 5 min. The pellet was resuspended in 650 µL water and divided equally between duplicate starch purification tubes as follows. Each subsample was carefully layered onto a 1-ml cushion of 80% (v/v) CsCl in a 1.5-ml tube and then centrifuged at 10,000 g for 5 min. The supernatants were discarded and the duplicate pellets were combined in 300 µl water. Starch pellets were each washed with 600 ul water, followed by one wash with 300 µl 2% SDS and three washes with 600 µl water. The final starch pellet was resuspended in 1 ml of water and frozen at -20°C prior to analysis. Duplicate 200-500-µl samples of each purified starch suspension were analyzed using a Multisizer 4e Coulter counter (Beckman Coulter) fitted with a 70-µm aperture tube and running in volumetric mode (counting 2 ml). The number of starch granules per endosperm, as well as the granule-size distributions were calculated. The morphology of the purified starch granules was examined using a Nova NanoSEM 450 (FEI, Hillsboro) scanning electron microscope.

**High-precision phenotyping of dry grains**

The shape and size of 400 dry grains (200 *LYS3* and 200 *lys3.c*) was captured and measured by customized Boxeed 2.1 (Labdeers) in 360° view every 10° resulting in 36 photos and table of measurements for every grain. Grain recognition and thickness measurements were performed on profile pictures of every grain using OpenCV library[36]. Thickness profiles were used for machine learning to define differences between *LYS3* and *lys3.c*. The trained model was further used for classification of $F_2$ grains from self-pollinated *lys3.c* heterozygotes.
