## Supplemental Materials for "Temporal genomic imprinting regulates grain filling in barley"

**Supplementary Figure 1 (Supports Figure 1).** Visualization of the candidate-imprinted and genotype-specific genes in F1 MOR-HS reciprocal crosses 8 and 12 days after pollination (DAP). Dots: pink/blue – maternally/paternally expressed genes (MEGs/PEGs, respectively); yellow/ green – genes with genotype (MOR/HS)-specific expression, respectively; grey – non-imprinted genotype independently expressed genes.

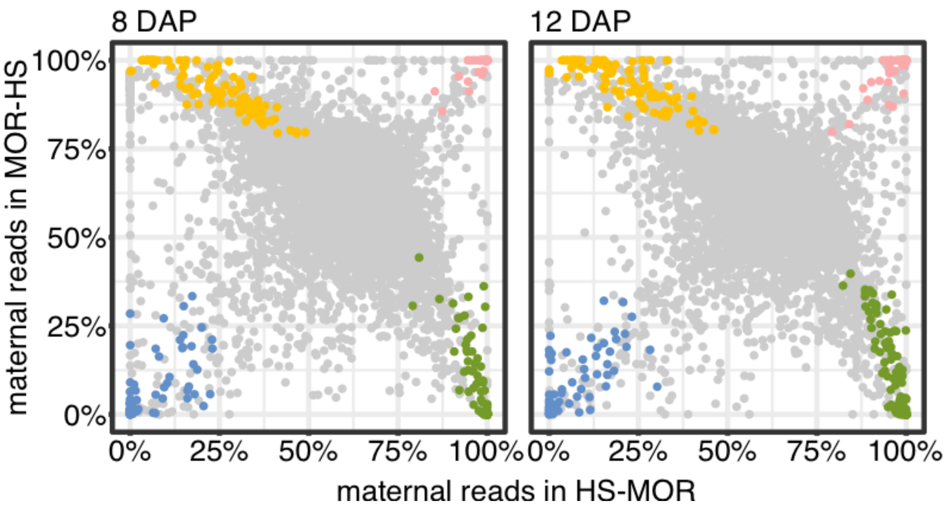

**Supplementary Figure 2 (Supports Figure 1).** Reverse transcription-PCR-based validation of the parent-of-origin specific expression of selected imprinted genes in barley. The information for every tested gene contains (top to bottom): NCBI Morex v3 gene identifier, genotype name - Morex (MOR), Ethiopian landrace (ETH) or wild barley (HS), corresponding genomic sequence with the diagnostic SNP (underlined), crossing combination (the maternal genotype is listed as the first) and corresponding cDNA sequencing trace files, and classification of the candidate as maternally expressed gene (MEG) or paternally expressed gene (PEG). Missing trace files indicate absence of diagnostic SNP in the given parental combination.

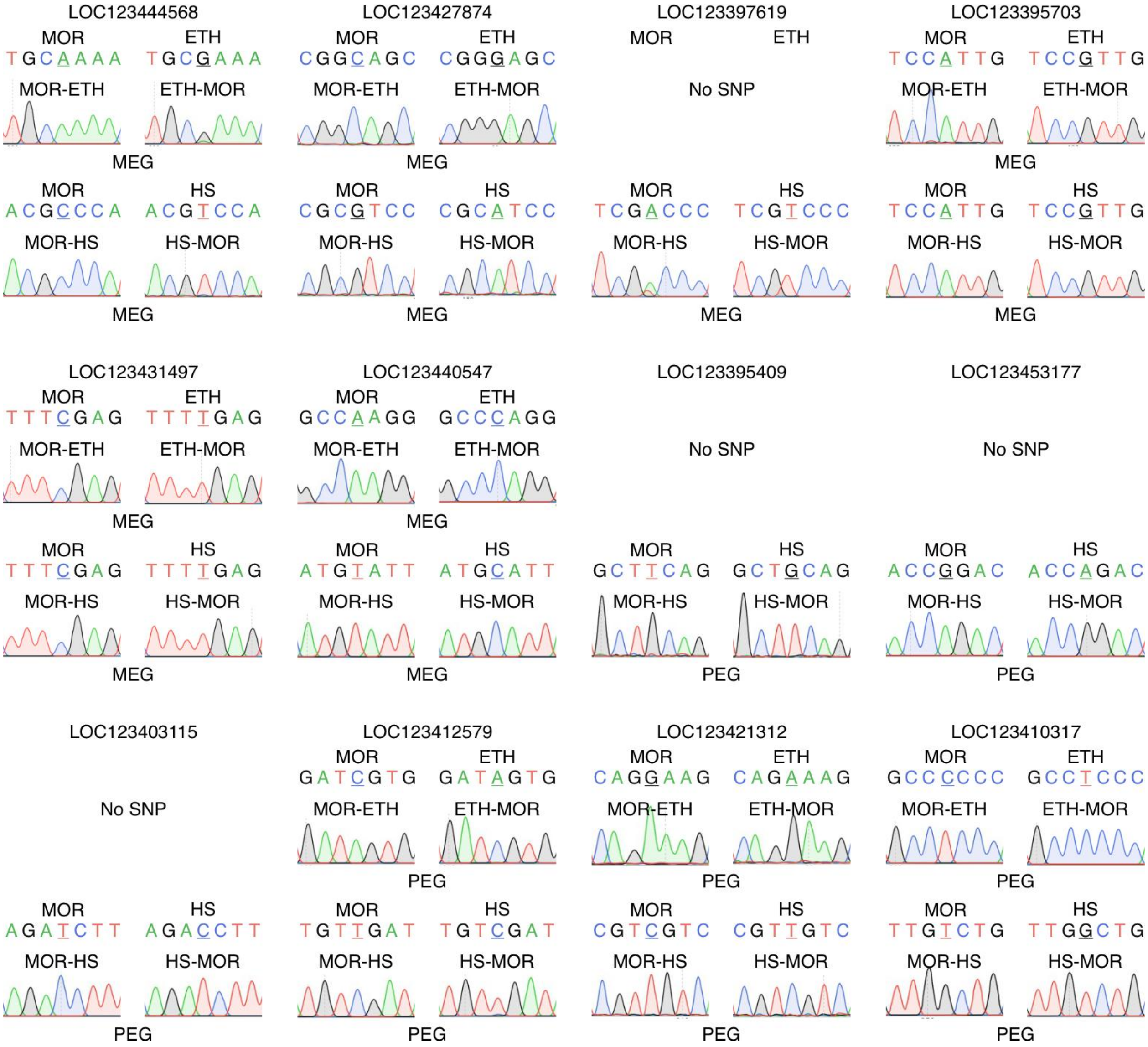

**Supplementary Figure 3 (Supports Figure 1)** Venn diagrams of the MEGs or PEGs at 8 and 12 days after pollination (DAP) endosperm in reciprocal crosses between Morex and Ethiopian landrace (MOR-ETH) or Morex and wild barley (MOR-HS).

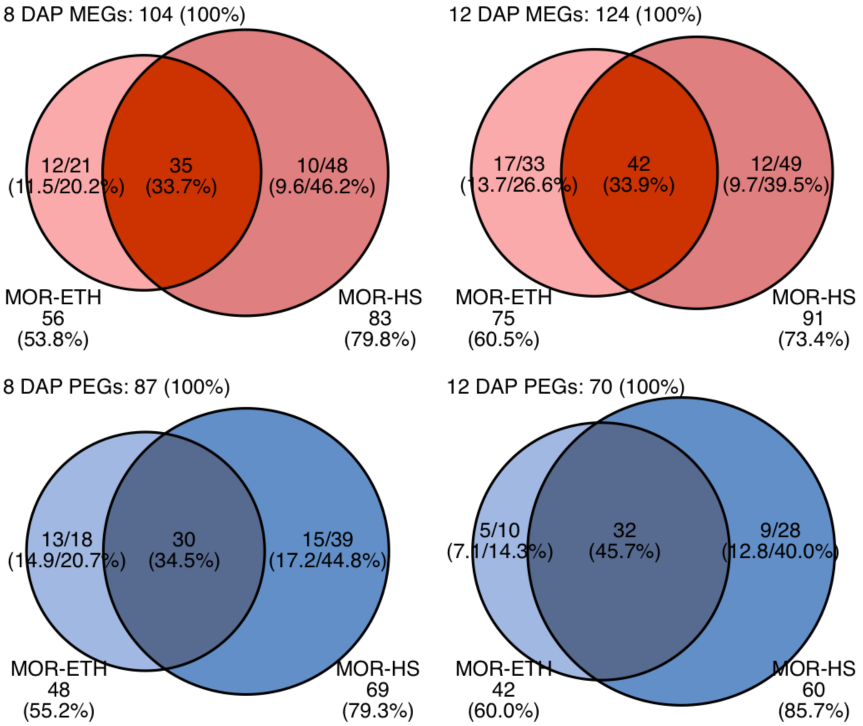

**Supplementary Figure 4 (Supports Figure 1)** Graphical representation of the KEGG pathways enriched for imprinted genes. (**a**) Ubiquitin-mediated proteolysis. (**b**) Plant hormone signal transduction pathway. Background color of proteins without and with NCBI Refseq annotation is white and green, respectively. The background of proteins coded by the maternally expressed genes (MEGs) is red and that of the paternally expressed genes (PEGs) is blue. Numbers below/above the marked gene indicate number of imprinted copies.

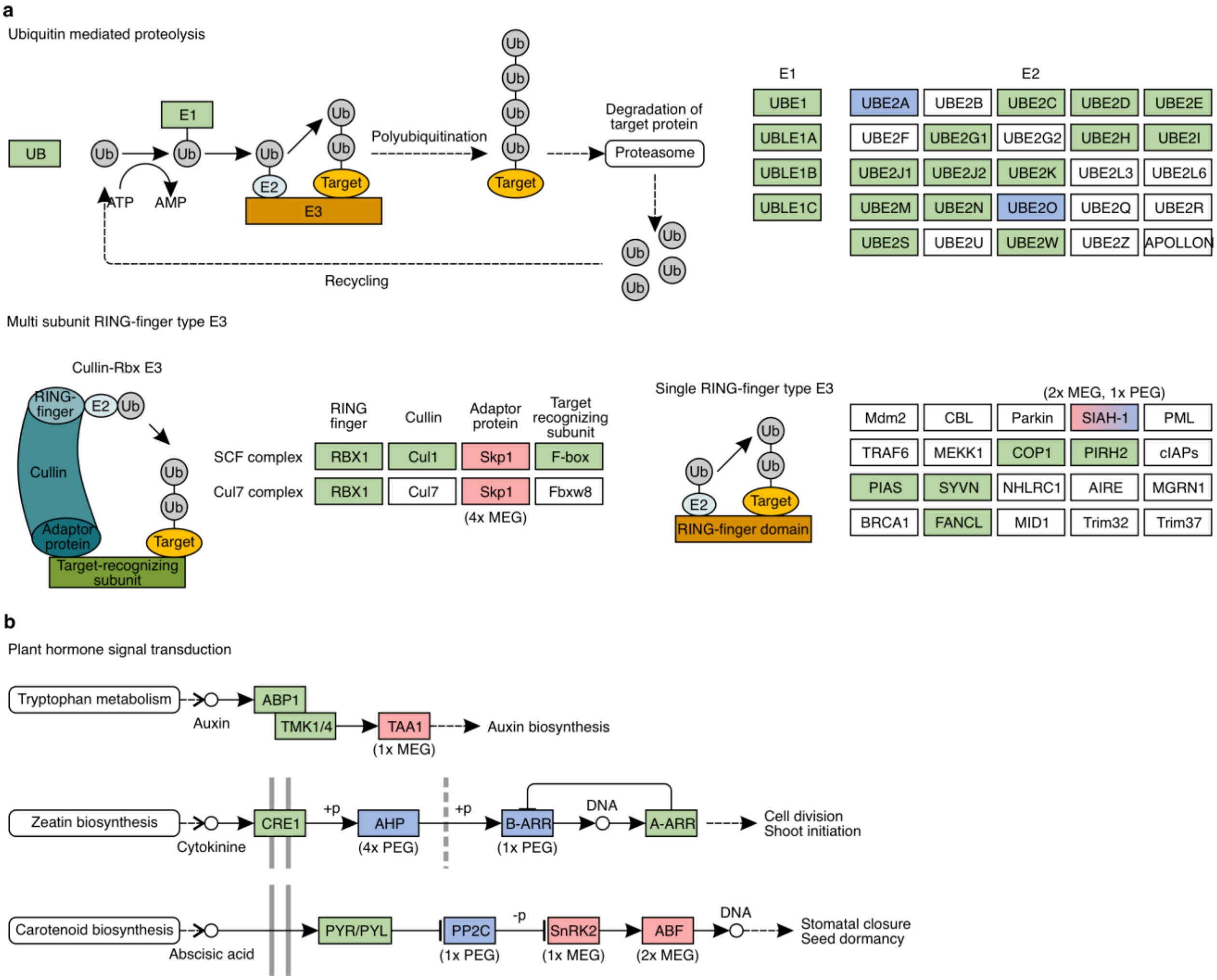

**Supplementary Figure 5 (Supports Figure 3)** Spatio-temporal dynamics of expression of individual barley imprinted genes from the k-means Cluster 1. Hierarchically clustered spatial transcriptomics bubble plots indicate expression of maternally expressed genes (MEGs) and paternally expressed genes (PEGs) at 6 and 12 days after pollination (DAP) grain sections. Circle sizes - percentage of expression; color - z-score expression; upper row – MEGs; lower row - PEGs. The x-axis represents individual tissues as indicated in the representative sections from Peirats-Llobet et. al, 2026.

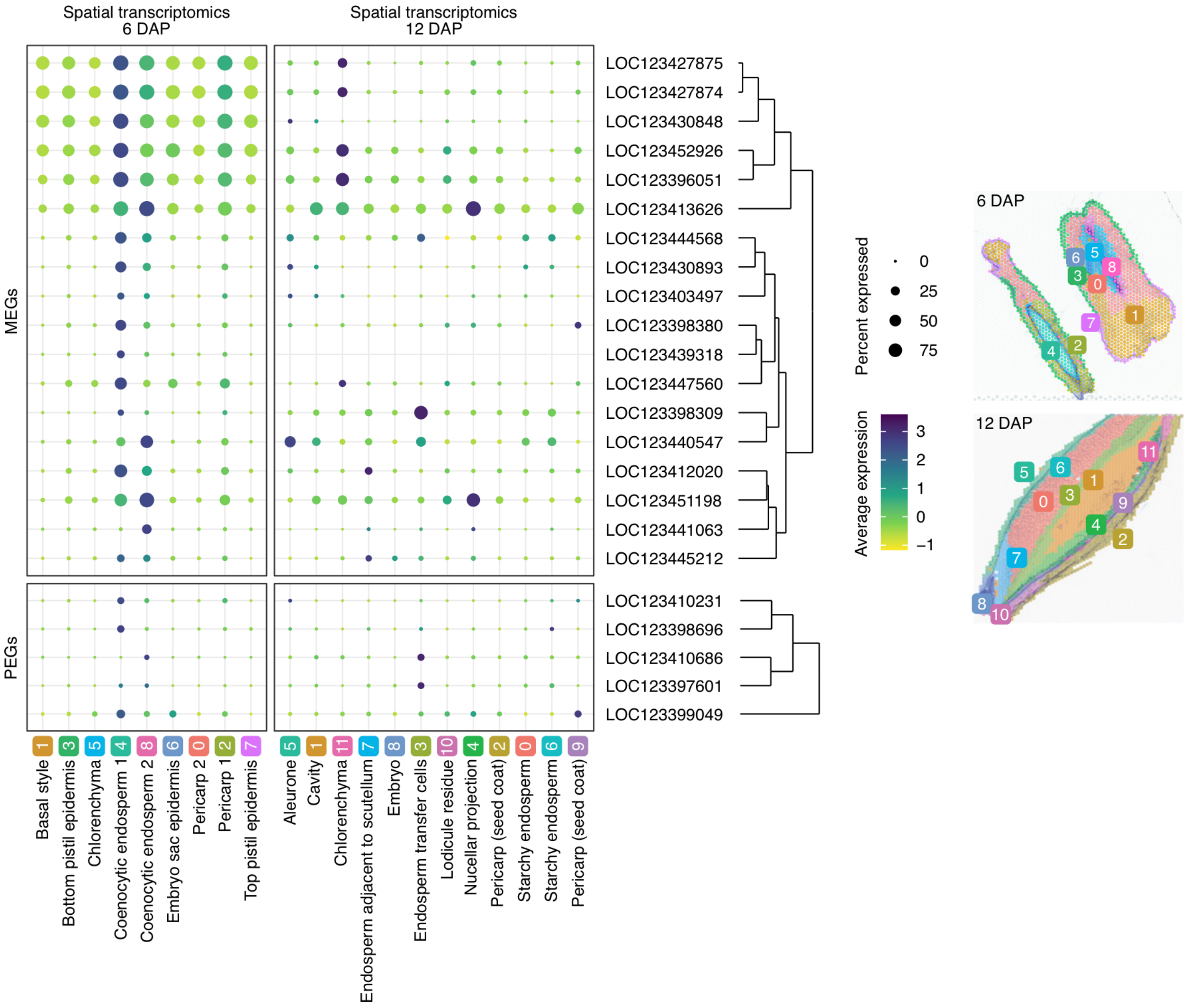

**Supplementary Figure 6 (Supports Figure 3)** Spatio-temporal dynamics of expression of individual barley imprinted genes from the k-means Cluster 2. Hierarchically clustered spatial transcriptomics bubble plots indicate expression of maternally expressed genes (MEGs) and paternally expressed genes (PEGs) at 6 and 12 days after pollination (DAP) grain sections. Circle sizes - percentage of expression; color - z-score expression; upper row – MEGs; lower row - PEGs. The x-axis represents individual tissues as indicated in the representative sections from Peirats- Llobet et. al, 2026.

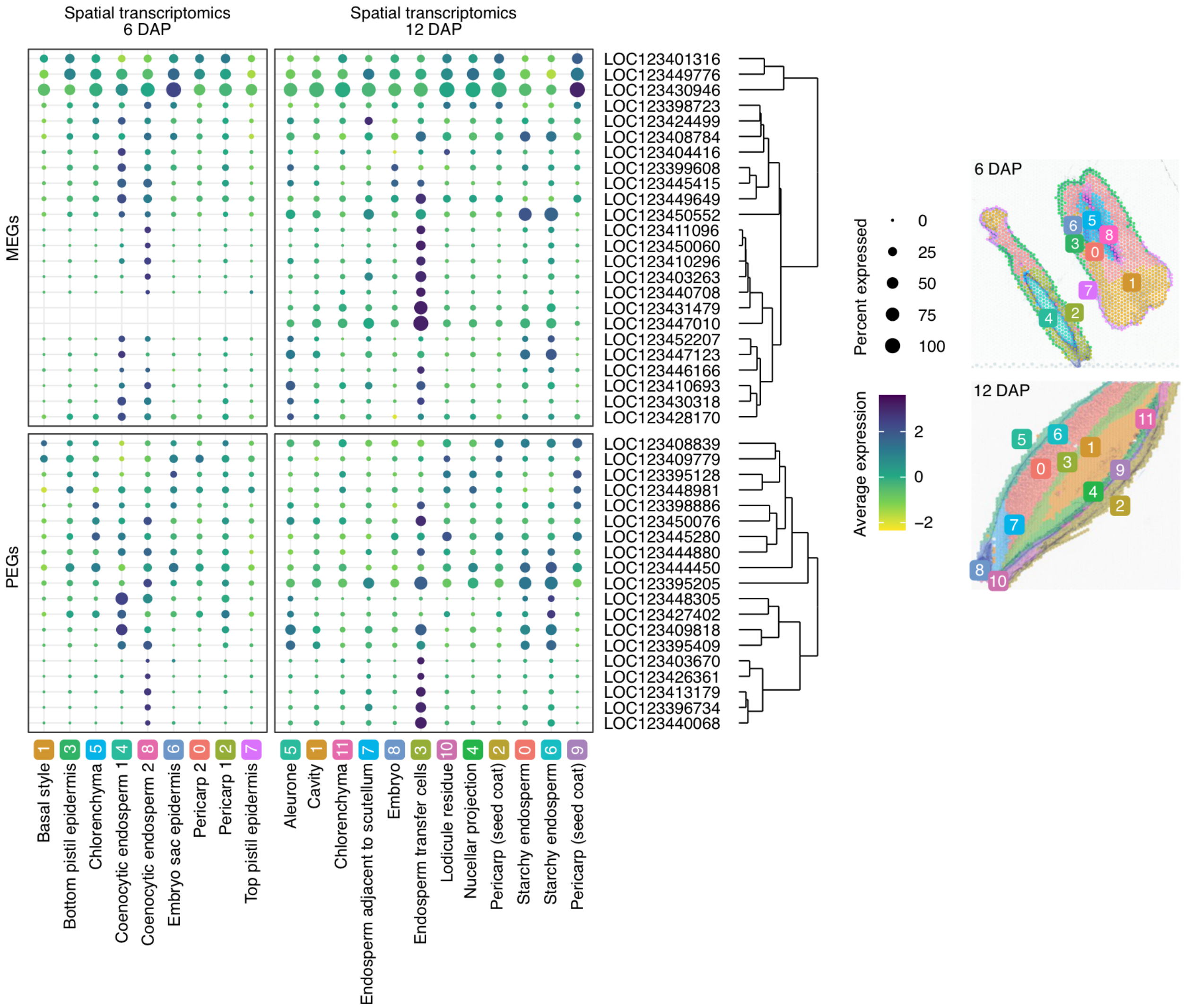

**Supplementary Figure 7 (Supports Figure 3)** Spatio-temporal dynamics of expression of individual barley imprinted genes from the k-means Cluster 3. Hierarchically clustered spatial transcriptomics bubble plots indicate expression of maternally expressed genes (MEGs) and paternally expressed genes (PEGs) at 6 and 12 days after pollination (DAP) grain sections. Circle sizes - percentage of expression; color - z-score expression; upper row – MEGs; lower row - PEGs. The x-axis represents individual tissues as indicated in the representative sections from Peirats- Llobet et. al, 2026.

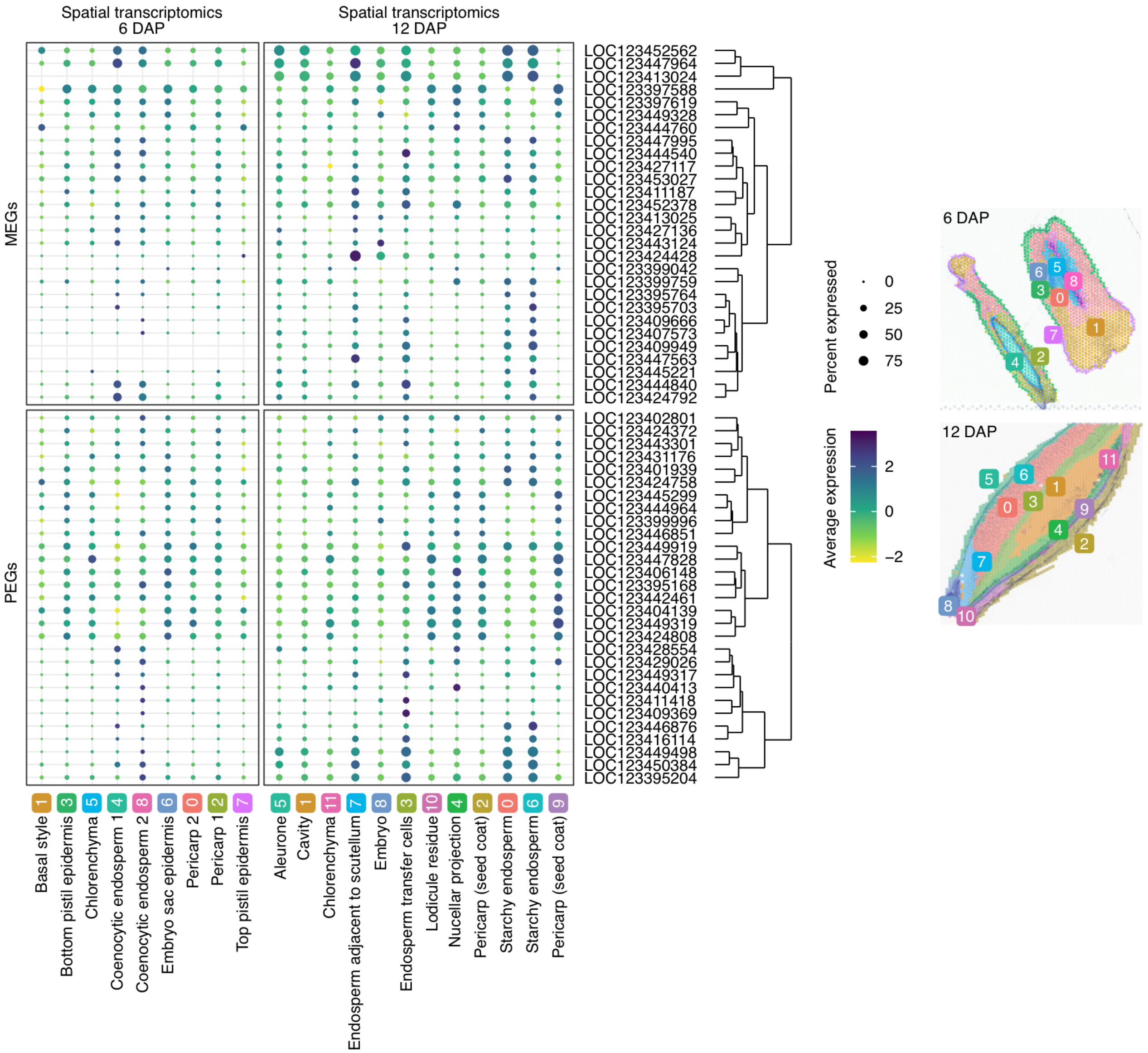

**Supplementary Figure 8 (Supports Figure 4).** Differentially expressed downregulated genes in reciprocal hybrid endosperms from wild-type Bomi (L) and *lys3.c* (l). Maternal genotype is always listed as the first one. (**a**) Venn diagram of significantly downregulated (FDR-adjusted p < 0.05, log2FoldChange < -1) genes in individual endosperms at 20 DAP compared to LL-L. (**b**) Differential expression of all genes in ll-L compared to LL-L endosperms at 12 DAP. Dots represent genes (Supplemental Data Set 11) without significant differential expression (grey), significantly down-regulated (light blue), significantly down-regulated at ll-l compared to LL-L at 12 and 20 DAP (dark blue), significantly up-regulated (salmon).

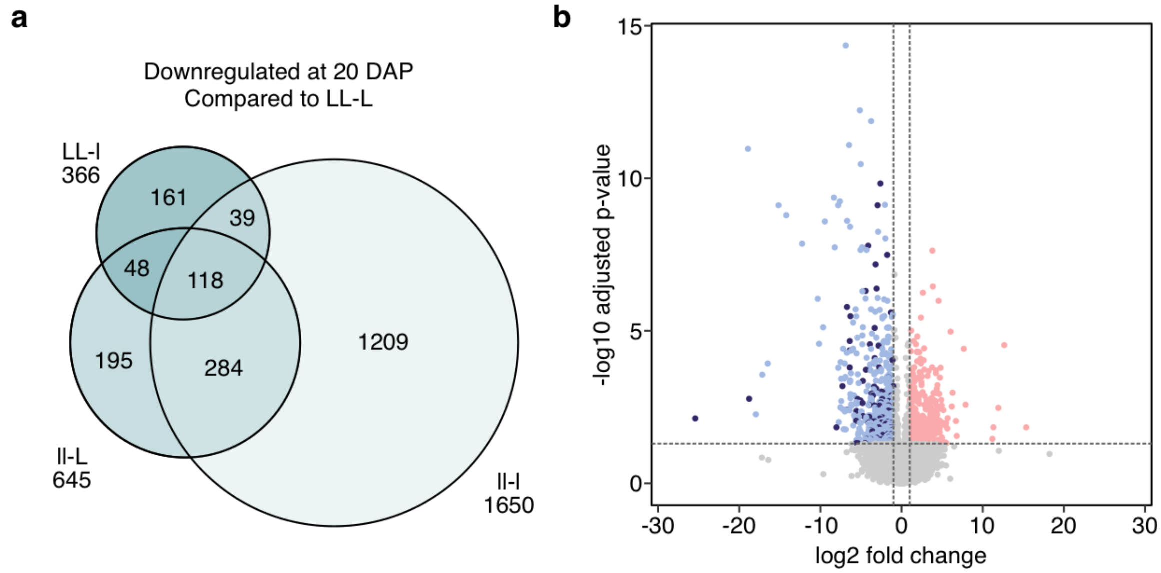

**Supplementary Figure 9 (Supports Figure 5)** Protein blots of HORDEINS in reciprocal wild-type *LYS3* (L) and *lys3.c* (l) crosses at given days after pollination (DAP). (**a**) Onset of HORDEIN accumulation in crosses using *lys3.c* allele. This blot is complemented by HORDEIN quantifications in blots at 16 and 20 DAP shown in Figure 5. (**b**) Accumulation of HORDEINS in crosses using *lys3.e* in RGT planet background at 16 and 20DAP. Equivalent fractions of total extracted proteins were loaded. (**c**) Difference in band intensities in individual crosses. Values marked with the same letter do not differ within the respective group according to two-way ANOVA with Tukey's HSD ($p < 0.05$).

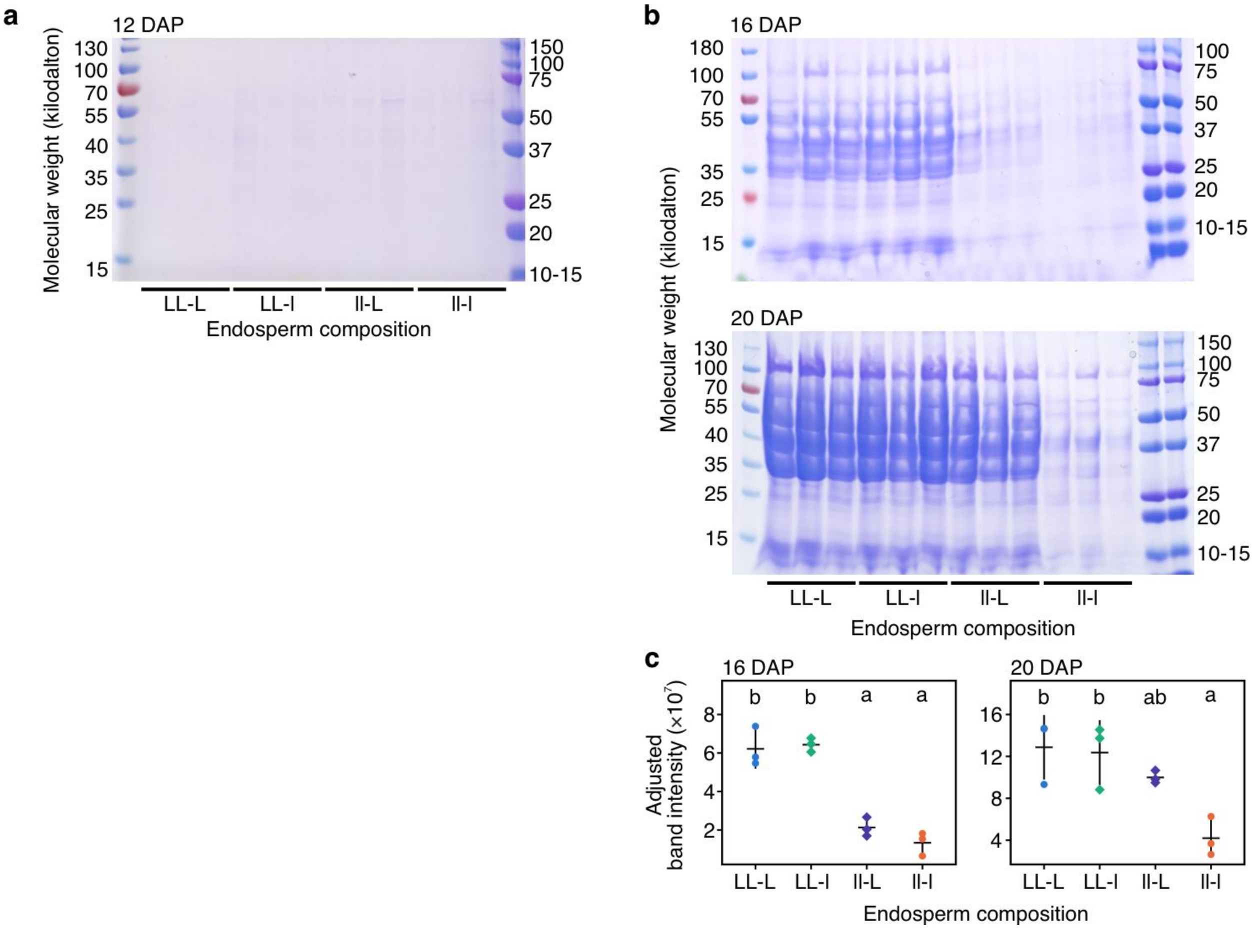

**Supplementary Figure 10 (Supports Figure 5)** Starch accumulation in reciprocal *LYS3* (L) and/or *lys3* (l) crosses. (**a**) Staining of starch granules (brown) using Lugol-based solution in transverse grain sections at given days after pollination (DAP). Scale bar applies to all images. (**b**) Histograms of size distribution of starch granules at 12 and 20 DAP. Differential volume represents the proportion (percentage) of the total volume of all measured particles that belongs to a specific particle size bin. The dashed lines indicate the mean size of the granules at 12 and 20 DAP in the wild type LL-L endosperm. (**c**) Quantification of starch granules per endosperm of individual *LYS3* and *lys3.e* crosses at given DAPs.

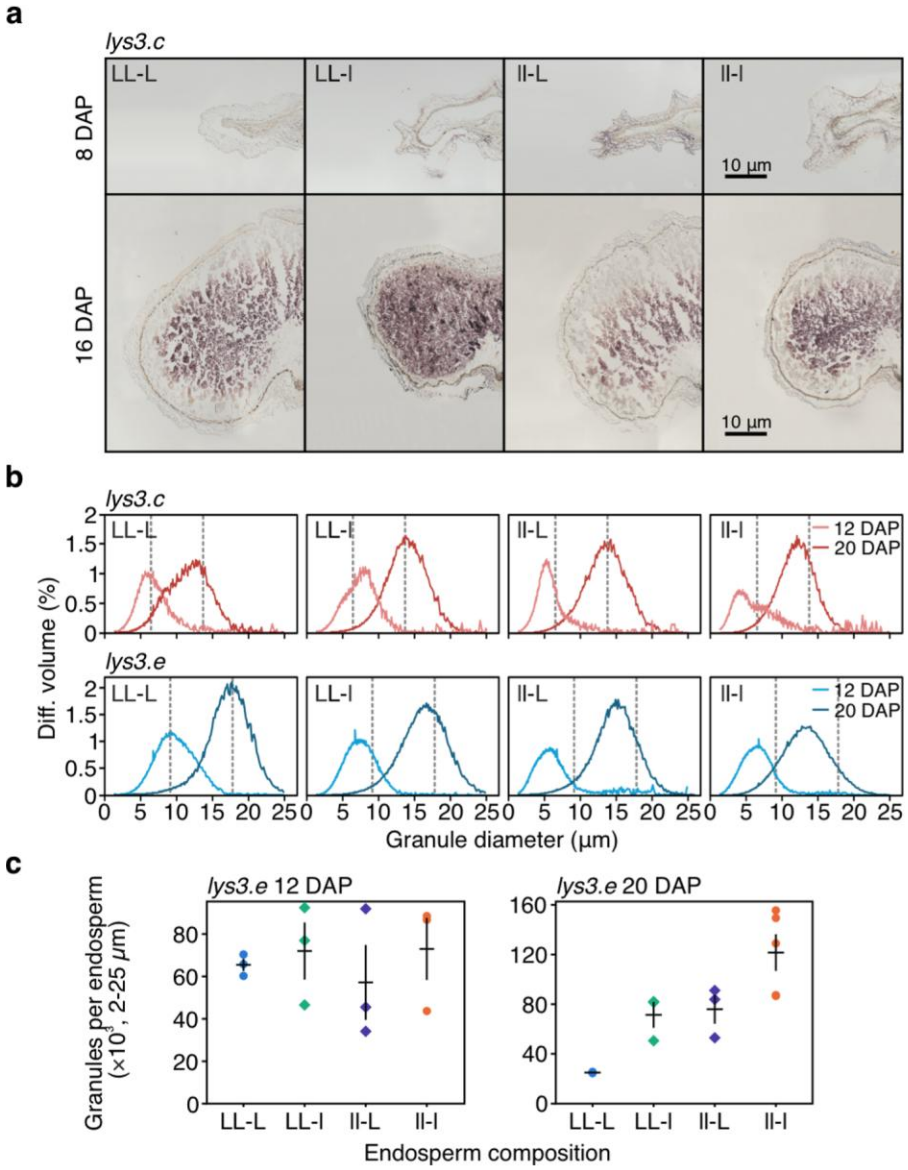

**Supplementary Figure 11 (Supports Figure 5)** High-precision phenotyping of dry grains of self-pollinated wild-type, hetero- and homozygous *lys3.c* plants. (**a**) Representative phenotype of *LYS3* and *lys3.c* mature dry grains. (**b**) Visualization of image processing pipeline showing segmentation, rotation of principal axis and thickness profile measurement. (**c**) Mean thickness profiles of 201 homozygous *LYS3* (blue) and 201 homozygous *lys3.c* grains (red). Lower panel shows difference in thickness between *LYS3* and *lys3.c* grains. Right panel shows cross-validated Linear discriminant analysis (LDA) scores. Histograms represent individual classifier score counts. Blue/red line represent fitted distribution for *LYS3*/*lys3.c*, respectively. (**d**) Two- (blue) and three-component (red) fit of Gaussian mixture model (GMM) to the heterozygote-delivered F2 seeds. (**e**) Simulation of 2000 synthetic F2 datasets under 3:1 segregation and comparing them to the observed distribution (red) with a Kolmogorov-Smirnov (KS) test.

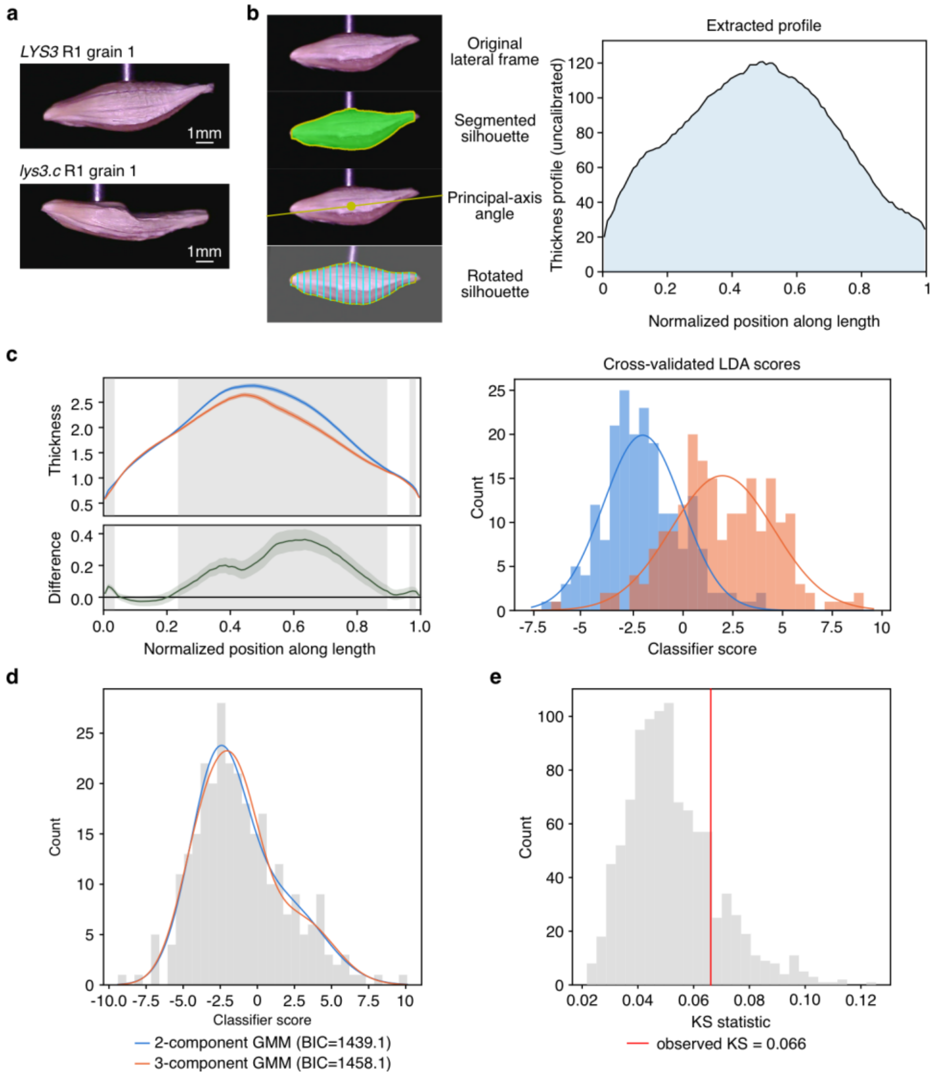

**Supplementary Figure 12 (Supports Figure 5)** Frequency of *LYS3* maternal and paternal reads. (**a,c**) Alignment of mapped reads at *LYS3* locus from reciprocal F1 MOR-ETH and MOR-HS crosses at given days after pollination (DAP). The reads were mapped on Morex reference genome and proportion of maternal and paternal reads is indicated by the proportion of individual single nucleotide polymorphisms. Read coverage (gray) with A, C, G and T SNPs in green, blue, orange and red, respectively. The blue gene models indicate whole predicted *LYS3* locus and its division into exons (thick rectangle), untranslated regions (thin rectangles) and intron (line with arrows). The gene is oriented from right (5') to left (3'). (**b,d**) Summarized percentage of maternal (salmon) and paternal (blue) *LYS3* RNA-seq reads from reciprocal F1 MOR-ETH and MOR-HS crosses endosperms at given DAP.

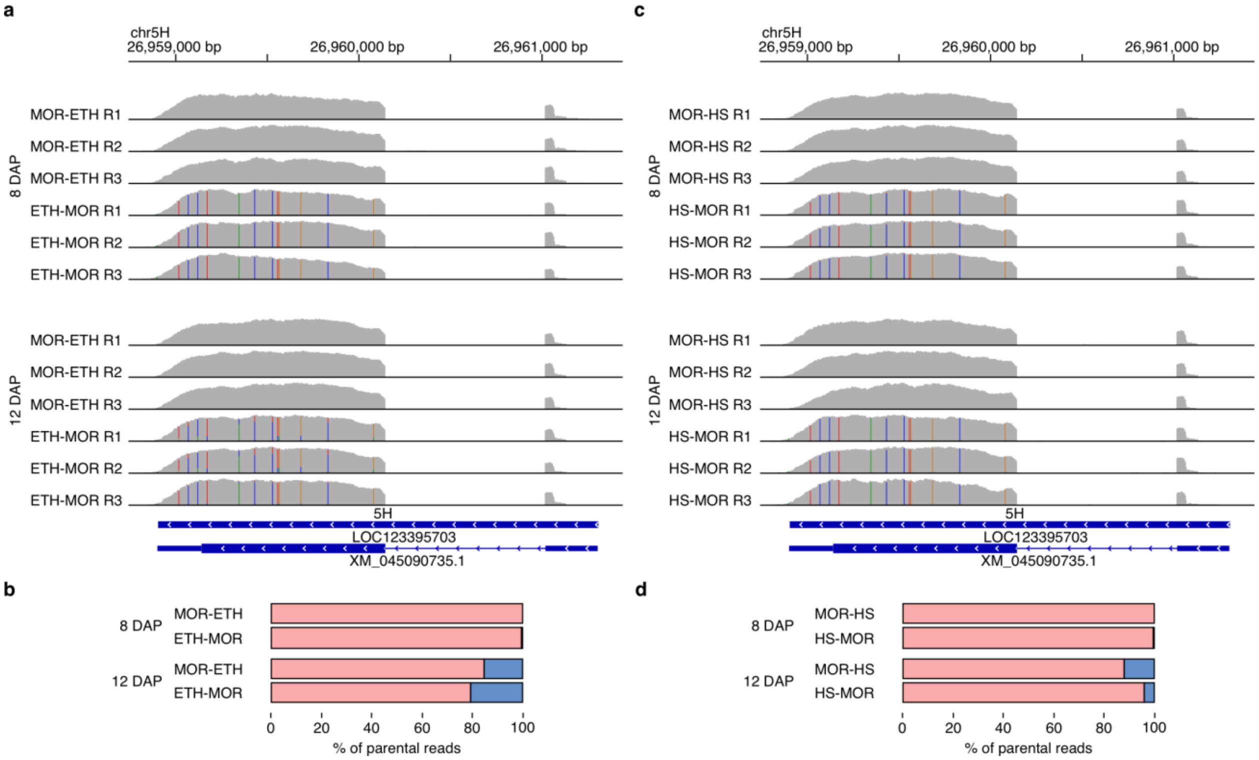

**Supplementary Figure 13.** Data analysis pipeline used for variant calling and identification of uniparentally expressed genes. Light blue boxes represent input/output files with file format indicated in grey. Arrows represents individual processing steps.

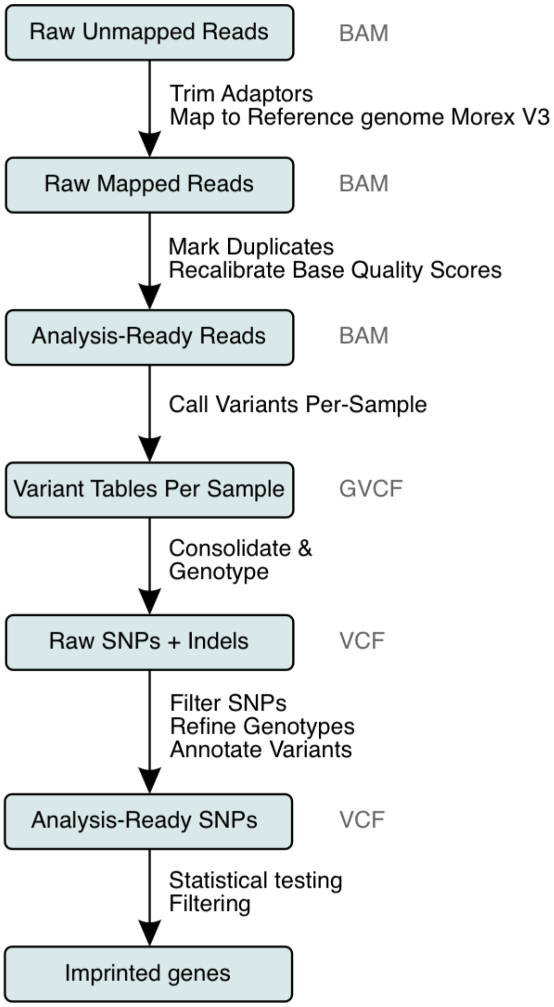

**Supplementary Table 1.** Primers used in this study

| Primer | Sequence (5'-3') | Gene | Application |
|---|---|---|---|
| lys3c_F1 | GCCGAAGCCAGACCAGAAGGTG | LOC123395703 | Bomi/*lys3.c* dCAPS |
| lys3c_R1 | GTCGGGAGGCCAGCAAAGTTCAT | LOC123395703 | Bomi/*lys3.c* dCAPS |
| lys3_FINDit48_F1 | TTTTCGTCCAACTCCAAGAGCA | LOC123395703 | RGT Planet/ *lys3.e* dCAPS |
| lys3_FINDit48_R1 | TCCAGAAATGAAGTCGTCCCA | LOC123395703 | RGT Planet/ *lys3.e* dCAPS |
| H2B_F1 | ATCCCAACGCAAATCTCCAACCTTC | LOC123440547 | Imprinting validation |
| H2B_R1 | TGGAGGTTTGTTTCAAGAAAATCGCCCAA | LOC123440547 | Imprinting validation |
| Agen_F1 | GAGCAATGGATGCCTCTGGATGCAA | LOC123395409 | Imprinting validation |
| Agen_R1 | TCAGGCCGTCAGAGCTTGTAAGATC | LOC123395409 | Imprinting validation |
| F-box_F1 | TGCAAGACCTGGAGGGGAATAGTCTC | LOC123453177 | Imprinting validation |
| F-box_R1 | TGAGATGTGAATCCAACGACTGGCT | LOC123453177 | Imprinting validation |
| Myb_F1 | ACTGGAATGCAATGCTGAGGGATACAC | LOC123403115 | Imprinting validation |
| Myb_R1 | TCAGTTGATCTGCAGTGGAATGGCTTG | LOC123403115 | Imprinting validation |
| Lys_N_F1 | TGGTACGACTTGCTGACGATCCTTCC | LOC123412579 | Imprinting validation |
| Lys_N_R1 | ACCATCAATGGAAGTCCGAGCACATC | LOC123412579 | Imprinting validation |
| Helix_F1 | GGTTTCTGTAGCAGGGGGTTCTTGAC | LOC123421312 | Imprinting validation |
| Helix_R1 | TTGCAGAAGTGATCGTCAKAGCTAGC | LOC123421312 | Imprinting validation |
| E3_F1 | GTCTCATGTAAGCACCCCGACAGA | LOC123410317 | Imprinting validation |
| E3_R1 | TTCGGATGTTACTGGCTTCGTCTTC | LOC123410317 | Imprinting validation |
| ALY3_F1 | CTGCTGAGGGCGCTGATGATTCG | LOC123397619 | Imprinting validation |
| ALY3_R1 | CCGACAGCGAGTACGGTCGAC | LOC123397619 | Imprinting validation |

| | | | |
|---|---|---|---|
| LTP_F1 | CACTACGTGCCACCATCGAGG | LOC123427874 | Imprinting validation |
| LTP_R1 | CTCTTGCGCGATTCCCATGAT | LOC123427874 | Imprinting validation |
| ULT1_F1 | AAGGGAGACTGCAGGAAAGTGGA | LOC123444568 | Imprinting validation |
| ULT1_R1 | GCTGTCAAATCTGCACATGTCCAG | LOC123444568 | Imprinting validation |
| NOT9A_F1 | AATCTACCCTGCACTTTCGCCT | LOC123431497 | Imprinting validation |
| NOT9A_R1 | GAAGGCTCATTGCCTATCTCCATG | LOC123431497 | Imprinting validation |
| LYS3_F1 | TGTTGTTGCATCGGACTTGG | LOC123395703 | Imprinting validation |
| LYS3_R1 | CTTTGCTAACGTCCTCCCGA | LOC123395703 | Imprinting validation |
| E3_exon2 | AATTCCGCCACAAACCCTAAGGCA | LOC123407184 | RT-qPCR control gene |
| E3_exon1 | CAGGAGCCCCCGTTGGATCA | LOC123407184 | RT-qPCR control gene |
| lys3_qPCR_ETH_MOR_F | TGAGAGGTGGTGCAGGAG | LOC123395703 | *LYS3* RT-qPCR |
| lys3_qPCR_ETH_R | ACATCCCAATGCTTGATCCG | LOC123395703 | *LYS3* RT-qPCR |
| lys3_qPCR_MOR_R | ACATCCCAATGCCTGATCCA | LOC123395703 | *LYS3* RT-qPCR |
